## Supplemental Figures for "Multi-omics characterization of partial chemical reprogramming reveals evidence of cell rejuvenation"

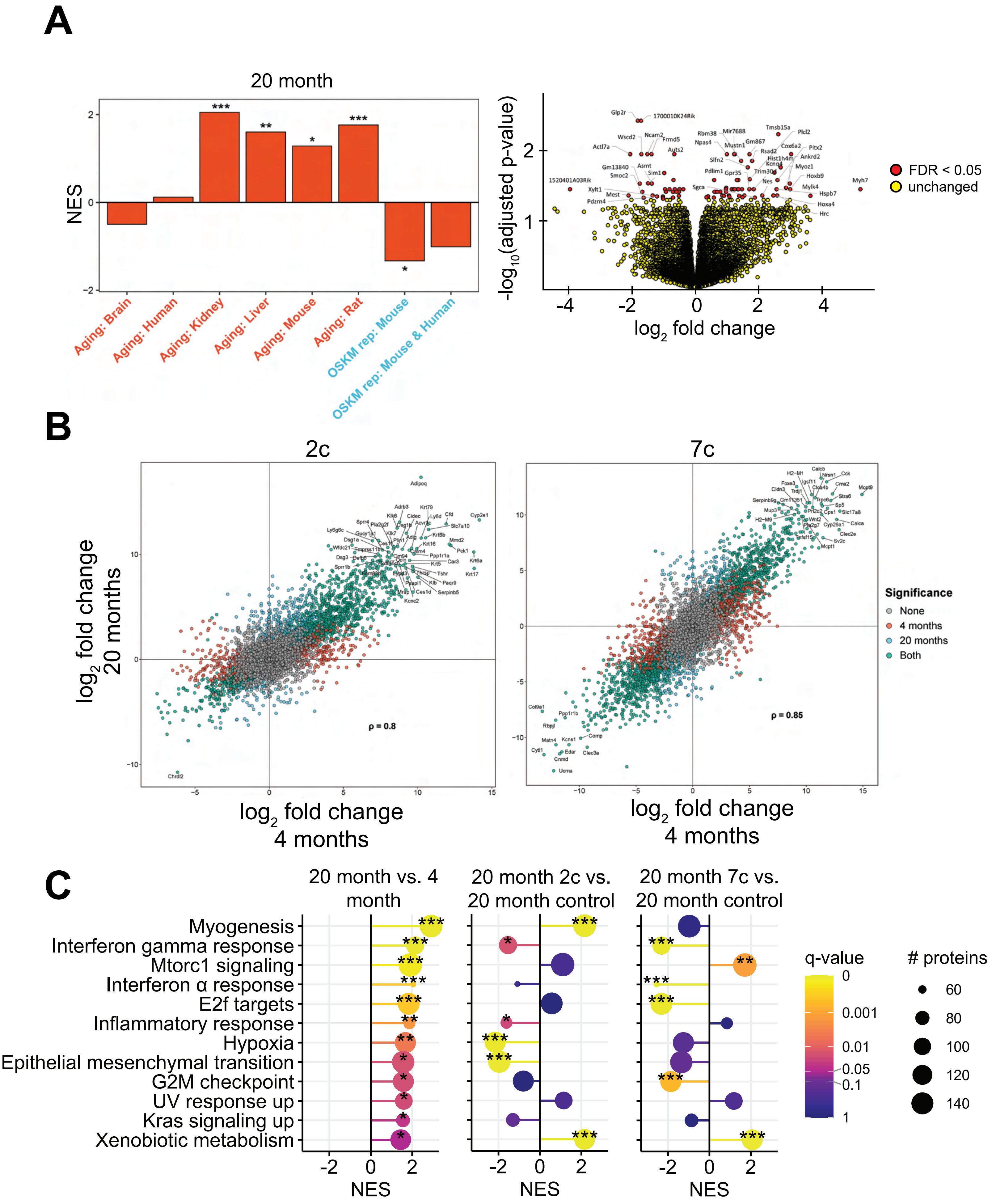


**Figure 2 - figure supplement 1**

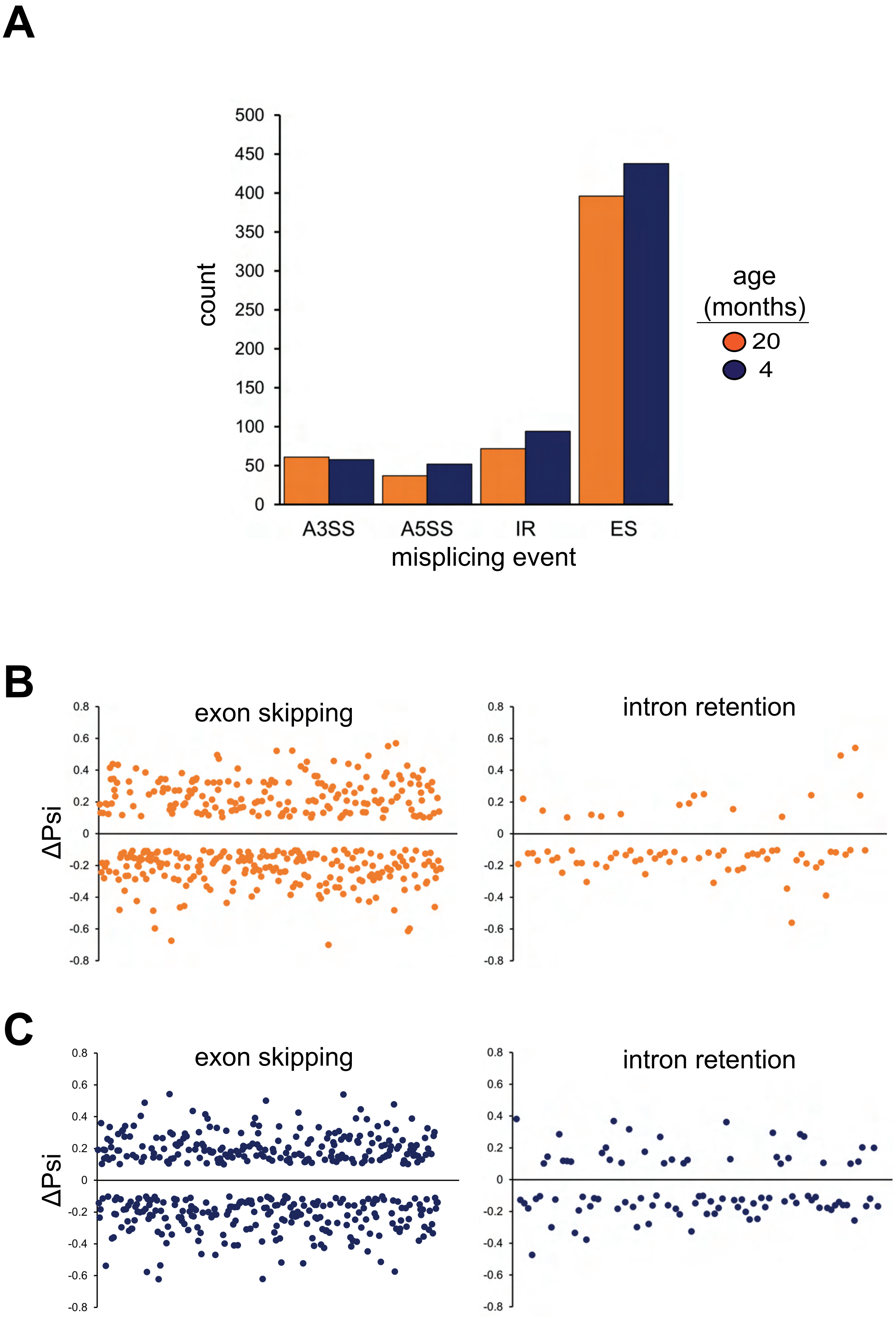


**Figure 2 - figure supplement 2**

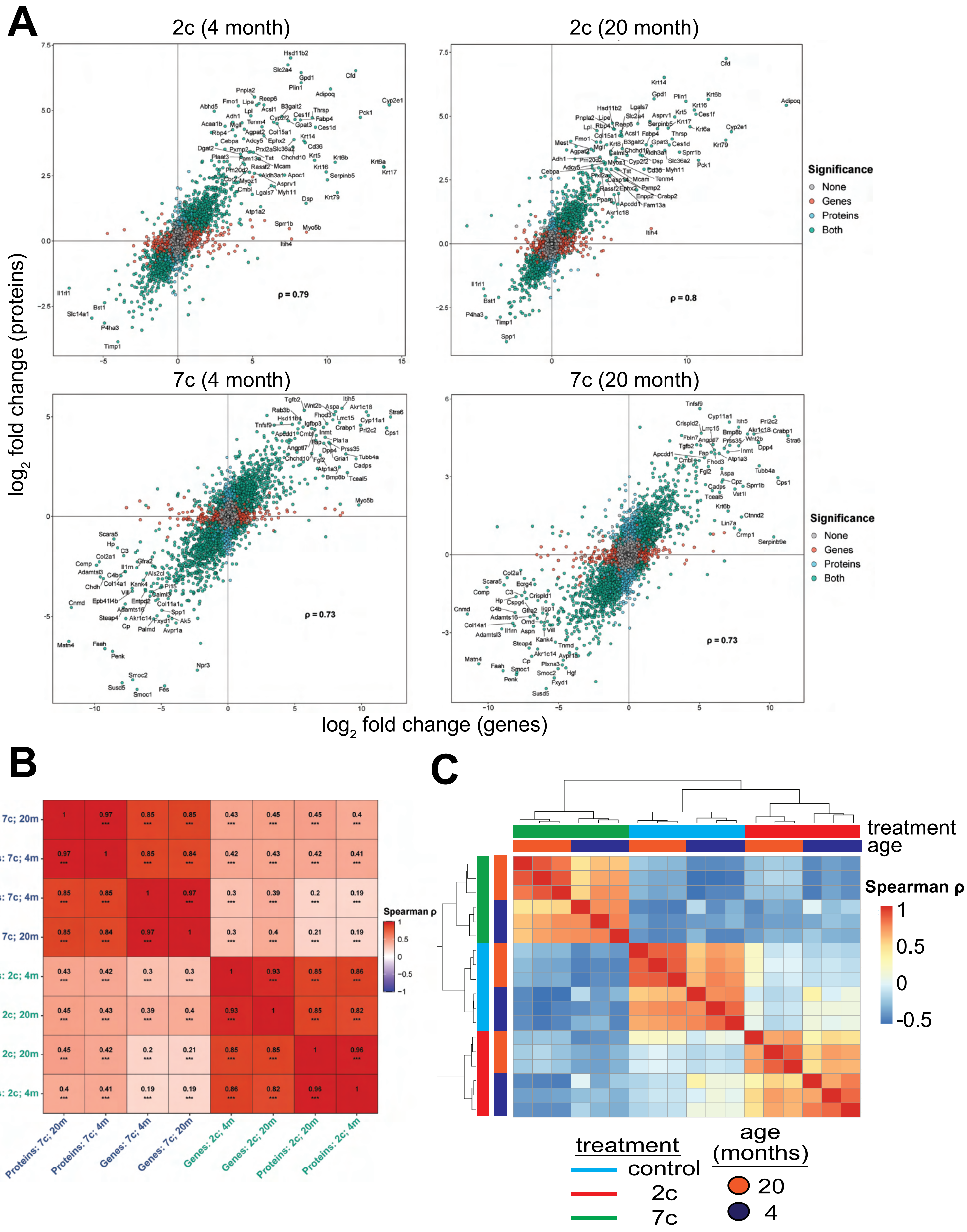


**Figure 3 - figure supplement 1**

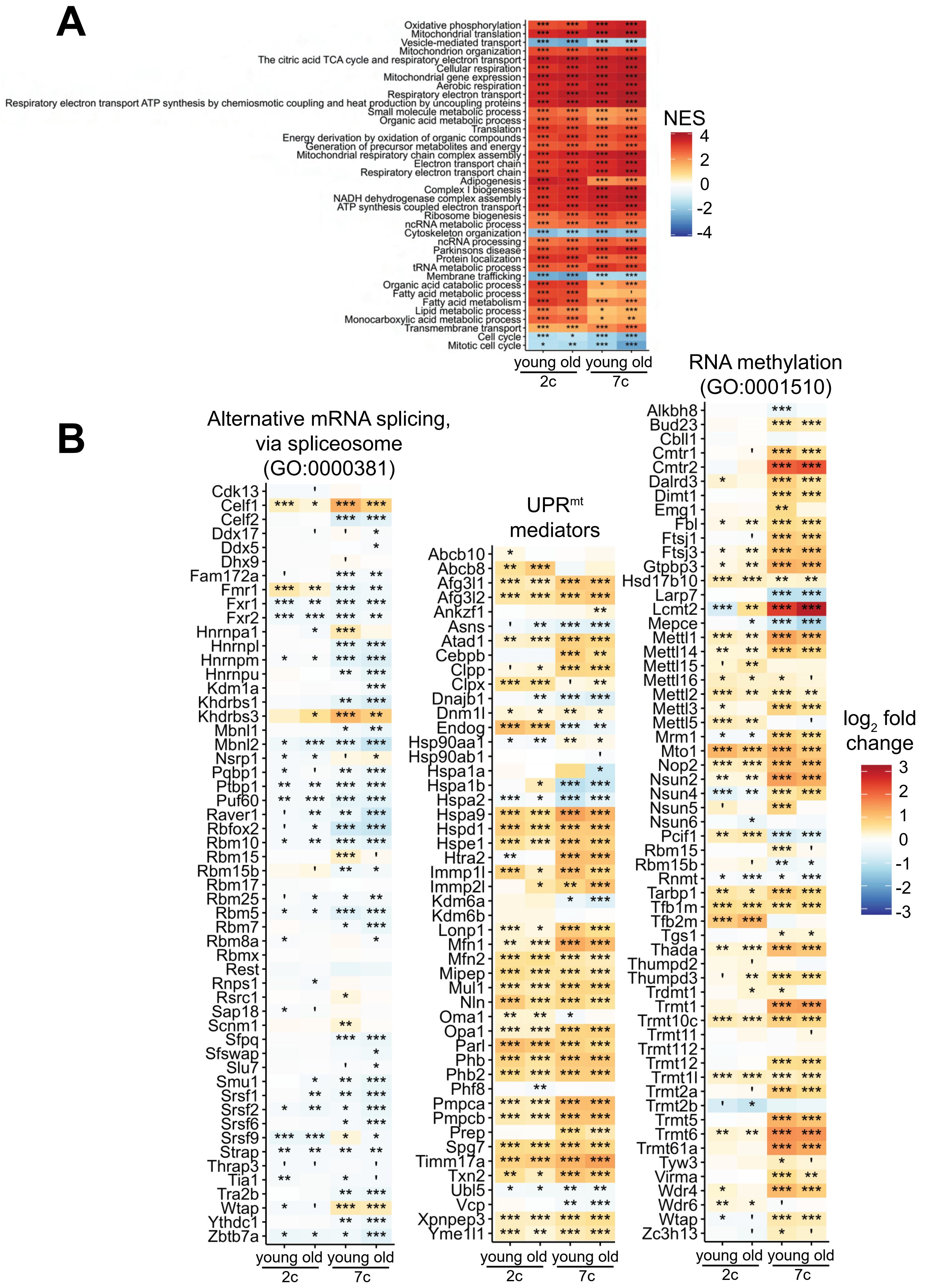


**Figure 3 - figure supplement 2**

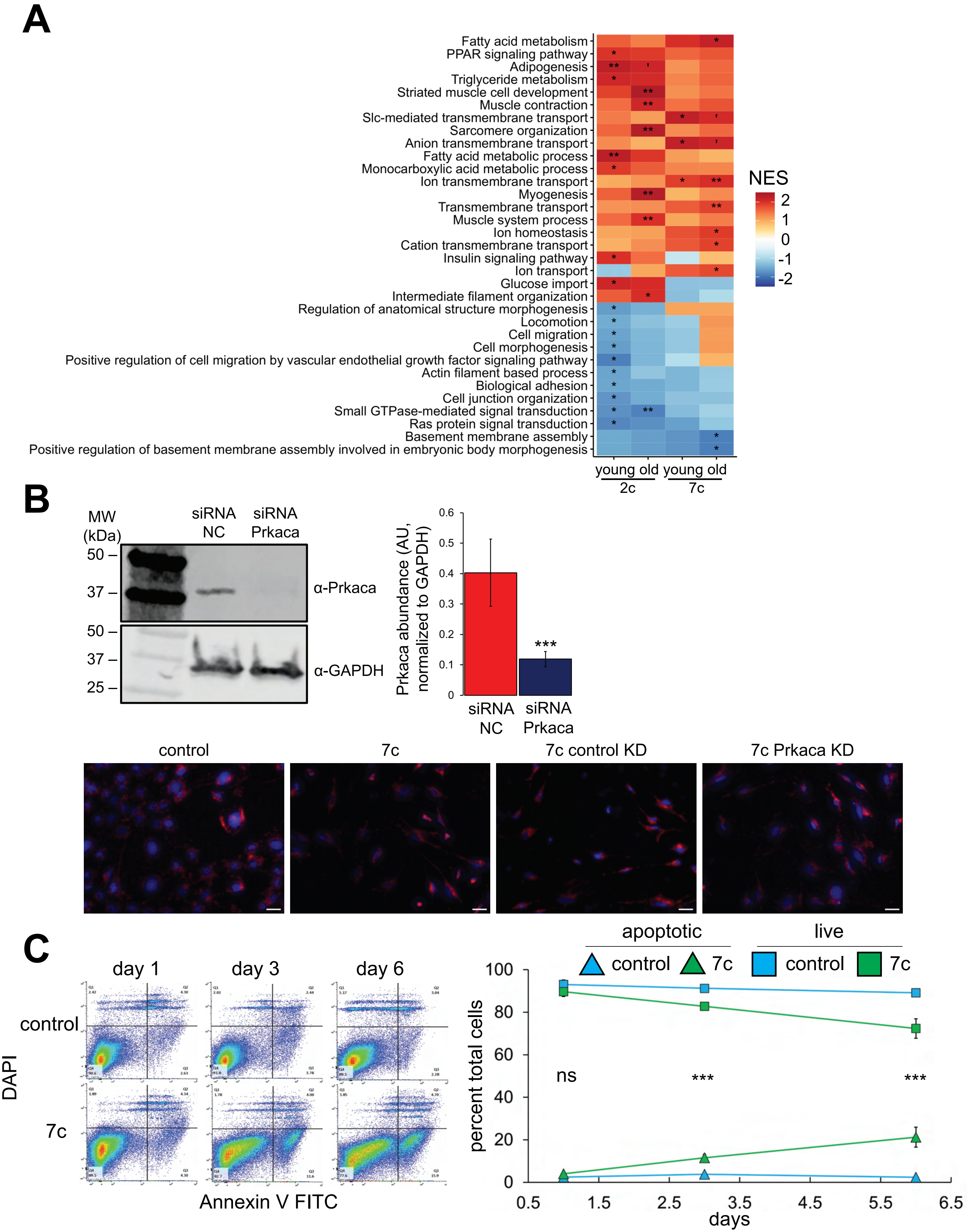


**Figure 5 - figure supplement 1**

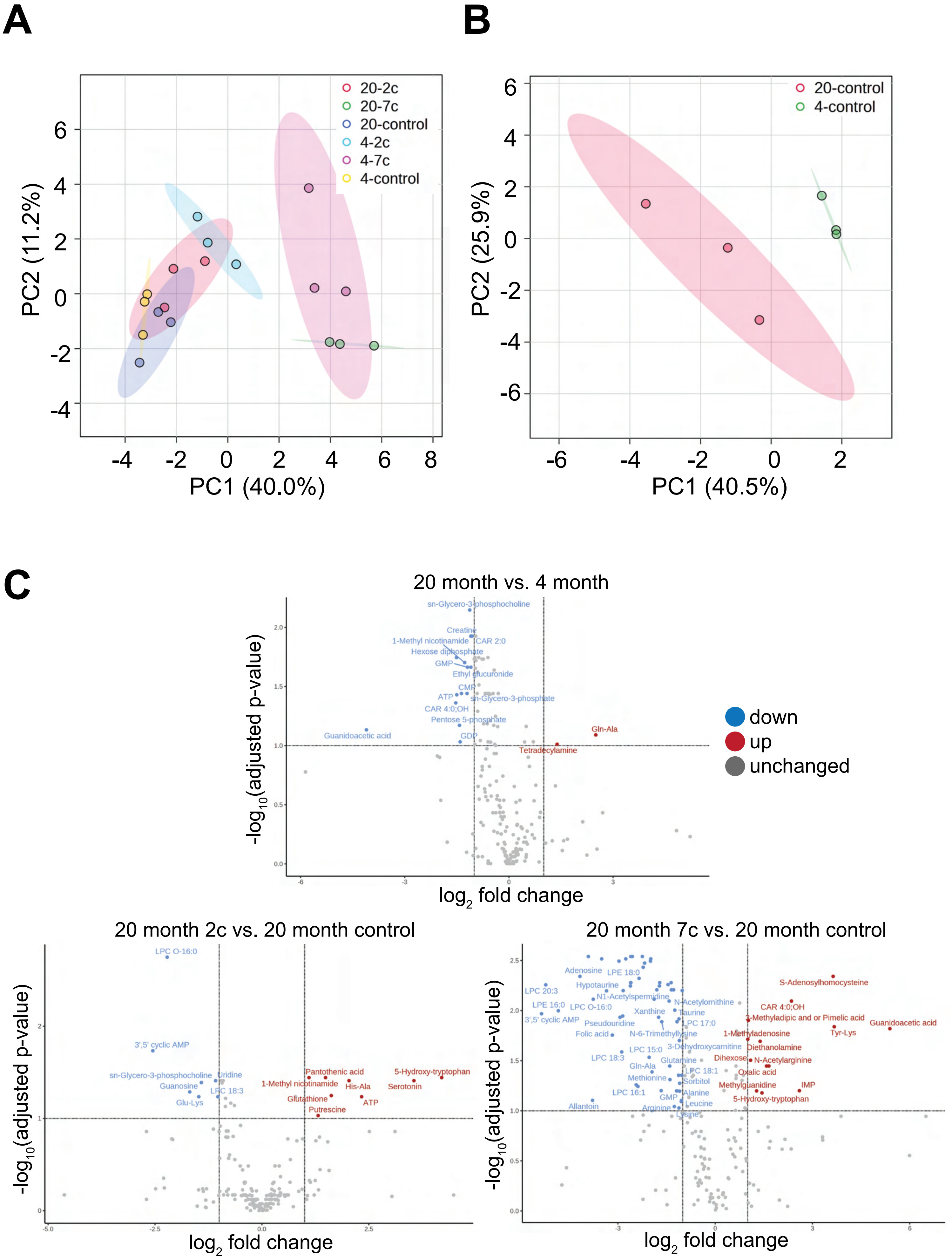


**Figure 6 - figure supplement 1**

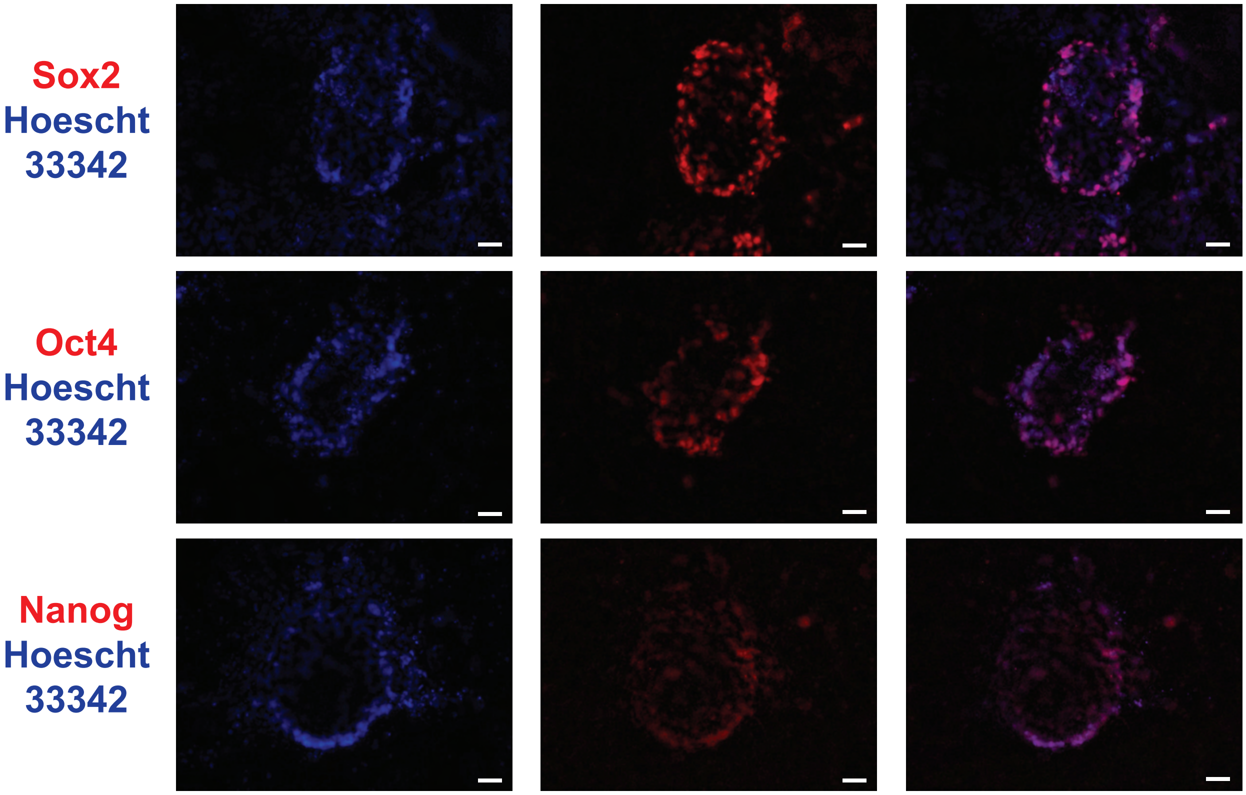


**Figure 7 - figure supplement 1**

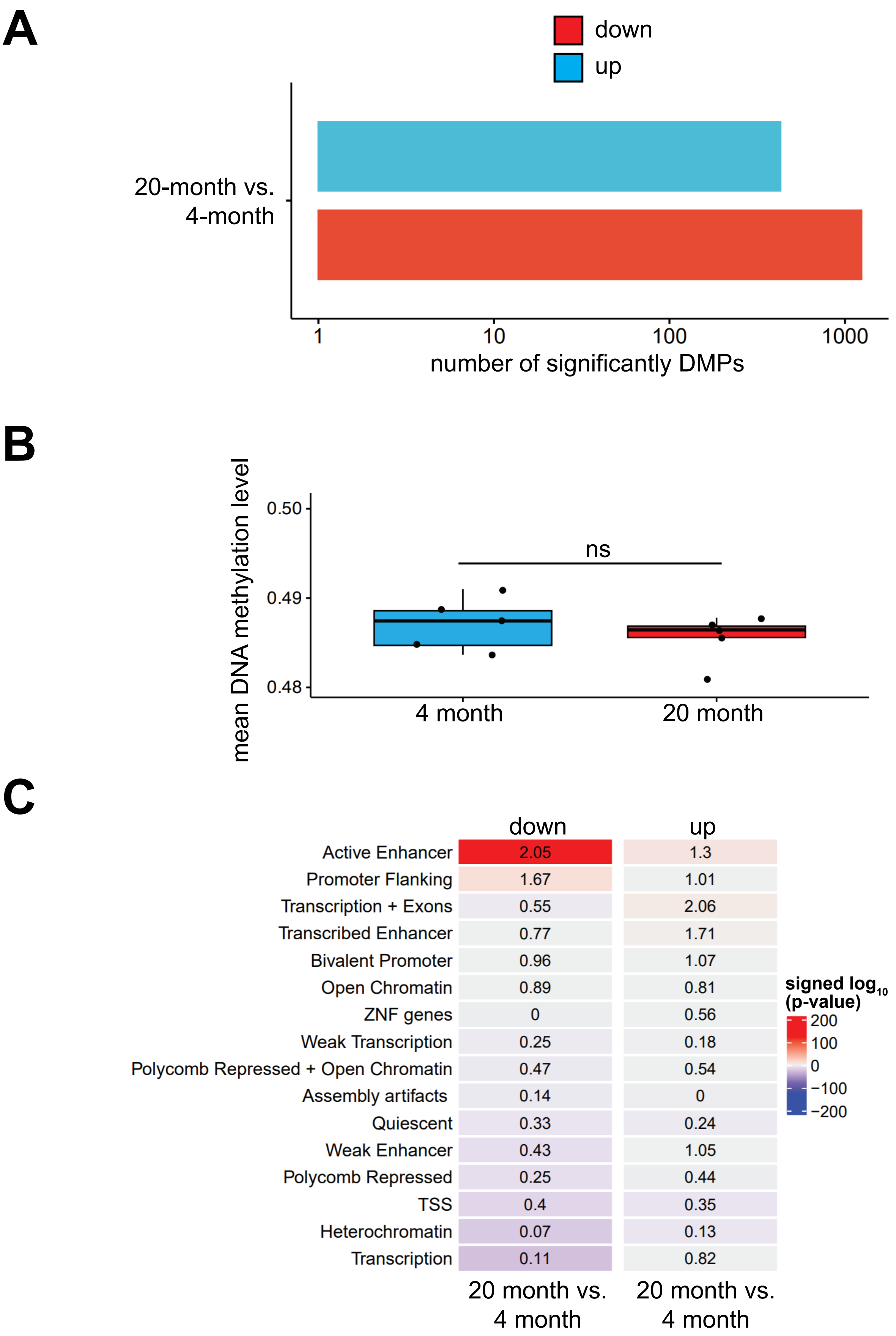


**Figure 7 - figure supplement 2**

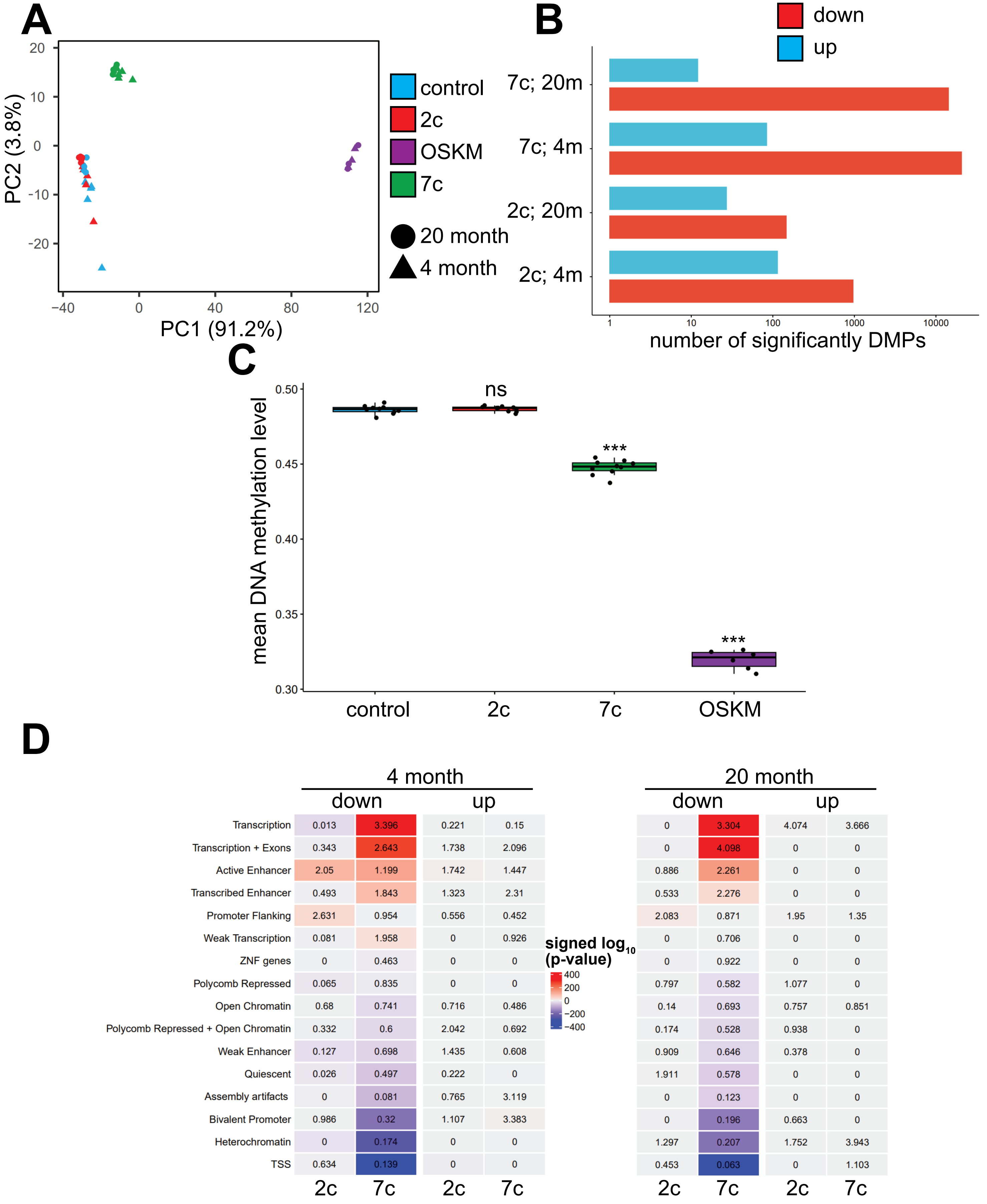


**Figure 7 - figure supplement 3**
